# The functional specificity of CDC42 isoforms is caused by their distinct subcellular localization

**DOI:** 10.1101/2023.02.27.528078

**Authors:** Yamini Ravichandran, Jan Hänisch, Kerren Murray, Vanessa Roca, Florent Dingli, Damarys Loew, Valentin Sabatet, Batiste Boëda, Theresia Stradal, Sandrine Etienne-Manneville

## Abstract

The small G-protein CDC42 is an evolutionary conserved polarity protein and a key regulator of numerous polarized cell functions, including directed cell migration. In vertebrates, alternative splicing gives rise to two CDC42 proteins: the ubiquitously expressed isoform (CDC42u) and the brain isoform (CDC42b), whose specific roles are not fully elucidated. The two isoforms only differ in their carboxy-terminal sequence, which includes the CAAX motif essential for CDC42 interaction with membrane. Here we show that these divergent sequences do not directly affect the range of CDC42’s potential binding partners, but indirectly influence CDC42-driven signaling by controlling the specific subcellular localization of the two isoforms. In astrocytes and neural precursors, which naturally express both variants, CDC42u is mainly cytosolic and associates with the leading-edge plasma membrane of migrating cells where it recruits the Par6-PKCζ complex to fulfill its polarity function. In contrast, CDC42b mainly localizes to intracellular membrane compartments, where it interacts with N-WASP. CDC42b does not participate in cell polarization but embodies the major isoform regulating endocytosis. Both CDC42 isoforms act in concert by contributing their specific functions to promote chemotaxis of neural precursors, demonstrating that the expression pattern of the two isoforms is decisive for the tissue-specific behavior of cells.

## Introduction

CDC42 is a small G protein of the Rho family (Etienne-Manneville and Hall, 2002) (Etienne-Manneville, 2004). Like most Rho GTPases, it switches between a GDP-bound inactive status and a GTP-bound form that can activate a large panel of effector proteins. Through these effectors RhoGTPases participate in numerous signaling cascades regulating a wide range of cellular responses. CDC42 interactions with its regulators and effectors generally occur on the cytosolic sides of cellular membranes whose planar geometry is thought to facilitate molecular interactions. Specific GEFs and GAPs regulate GTP-GDP cycling of CDC42 (Hodge and Ridley, 2016). The membrane localization of CDC42, like of other RhoGTPases, relies on the post-translational prenylation of the cysteine included in the carboxy-terminal CAAX box. This modification promotes the protein anchorage in the lipid bilayer. GDIs can interact with the lipid anchors of Rho GTPases to sequester them in the cytosol and thereby inhibit their functions (Hodge and Ridley, 2016; Johnson et al., 2009; Nomanbhoy et al., 1999).

The Rho GTPase CDC42 is more particularly known for its fundamental role in the control of polarity during cell asymmetric division, cell differentiation and cell migration in organisms ranging from yeast to mammals (Etienne-Manneville, 2004). CDC42–mediated signaling controls cytoskeleton rearrangement, which affects various actin and/or microtubule-dependent cellular processes and plays a key role in clathrin-dependent and clathrin-independent endocytosis, in exocytosis and, in vesicle transport (Chadda et al., 2007; Chen et al., 2005; Erickson and Cerione, 2001; Harris and Tepass, 2010; Mayor et al., 2014; Ridley, 2006; Sabharanjak et al., 2002; Wu et al., 2000). Functional dysregulation of CDC42 has been implicated in the pathology of several disease states and developmental disorders, including cancer (Aspenstrom, 2018; Martinelli et al., 2018). Studies using constitutively active or dominant-negative CDC42 mutants showed that CDC42 acts an oncoprotein which promotes cellular transformation and metastasis in the context of loss of polarity (Fidyk et al., 2006; Haga and Ridley, 2016; Johnson et al., 2010; Reymond et al., 2012). On the contrary, gene knockout studies suggested that CDC42 might also function as a tumor suppressor, because targeted knockout of the *CDC42* gene in hepatocytes or blood stem/progenitor cells resulted in the development of hepatocellular carcinoma or myeloproliferative disease in mice (van Hengel et al., 2008). This ambiguity makes understanding the role of CDC42 driven polarization in the context of cancer biology challenging.

Previous CDC42 studies have been mostly focused on the ubiquitous CDC42 isoform (CDC42u). However, the human *CDC42* gene located on chromosome 1 gives rise to three transcripts via alternative splicing, which translate into two distinct CDC42 isoforms. CDC42u, initially described as placental CDC42, does not include the exon 6 but an alternative exon 7 and is ubiquitously expressed (Marks and Kwiatkowski, 1996). In contrast, the so-called brain isoform (CDC42b), is generated by translation of the exons 1-6 and was initially detected in brain tissue. The expression of the ubiquitous isoform CDC42u in neuronal precursors and non-neuronal cells switches to a stable co-expression of the CDC42b and CDC42u isoforms at the single-neuron level. During neurogenesis, CDC42u specifically drives the formation of neuroprogenitor cells, whereas CDC42b is essential for promoting the transition of neuroprogenitor cells to neurons. However, the molecular mechanisms responsible for the specific functions of these two isoforms and their role in non-neuronal cells remain unknown. In fact, both isoforms were found to be expressed in a range of commonly used laboratory cell lines, including HEK and MDCKII cells (Wirth et al., 2013). This splice variant may thus also be expressed in non-brain tissue cells underlining the need to clarify the functional differences between the two CDC42 variants in non-cell specific functions. By inactivating specifically each of the two CDC42 variants in brain cells such as astrocytes and neural precursor cells (NPC), we unravel the specific localization and functions of the two isoforms during persistent directed migration.

## Results

### Cell polarization relies on ubiquitous CDC42

To determine whether CDC42 isoforms had different functions, we used primary astrocytes, glial cells in which we determined the expression levels of CDC42 isoforms. Primary rat astrocytes were transfected with siRNAs designed to selectively knockdown brain (si-b1, si-b2) or ubiquitous (si-u1, si-u2) CDC42 or with a siRNA targeting a common sequence (si-both) to simultaneously inhibit both isoforms. We developed a CDC42b specific antibody to assess the specific expression of this isoform (Fig. S1A, S1B). Unfortunately, we could not obtain an antibody specific for CDC42u. Thus, we used qPCR to quantify CDC42u and total CDC42 mRNA level. Knockdown of each isoform revealed that roughly 15-20% of total CDC42 mRNA in astrocytes encodes the brain isoform (Fig. S1C, S1D) (Yap et al., 2016).

We then chose to determine the contribution of each CDC42 isoform in cell directed migration, a more complex and integrated cell function. Control and CDC42 knockdown cells were submitted to a scratch-induced migration assay, a well-characterized model system in which we have previously demonstrated that CDC42 is activated by an integrin-mediated signaling and involved in microtubule-dependent front-rear cell polarization, Golgi and centrosome reorientation, and directed membrane recycling (Etienne-Manneville and Hall, 2001; Osmani et al., 2006). Knockdown of both CDC42 isoforms did not influence the migration velocity, but led to a strong decrease in directionality (83% to about 55%) and persistence (85% to about 60%) of migration (Fig. 1A-1C). The specific depletion of CDC42u led to similar results with a significant decrease in the directionality and persistence of migration, whereas CDC42b-depleted cells migrated similarly as control astrocytes (Fig. 1A-1C).

**Figure 1:**
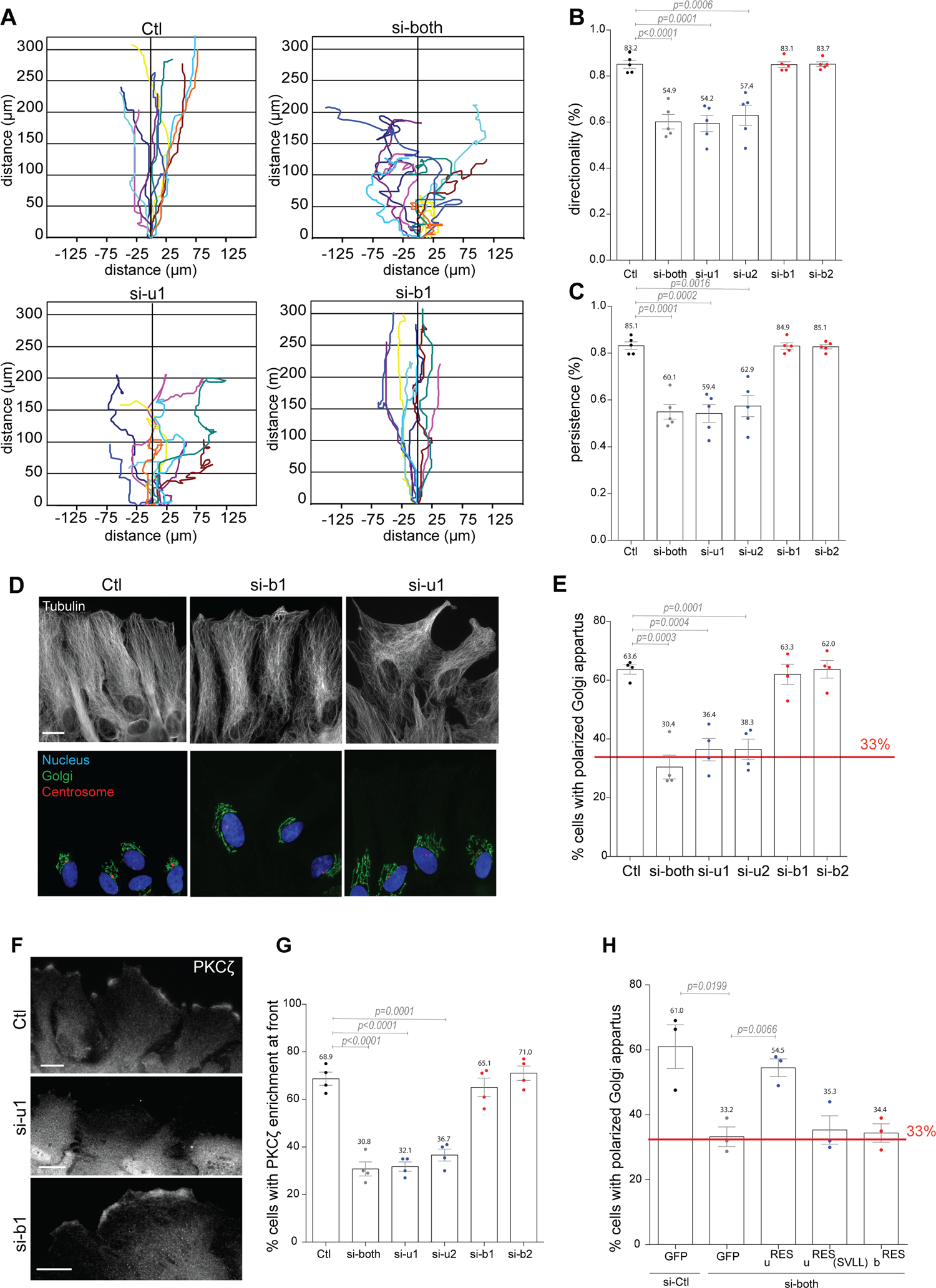
CDC42u, but not CDC42b control cell polarization and directed and persistent migration of astrocytes. **(A)** Representative trajectories of astrocytes transfected with siRNAs against the ubiquitous (si-u1), the brain (si-b1) or both (si-both) CDC42 isoforms and migrating in an in vitro wound healing assays for 16h. Directionality **(B)** and directional persistence **(C)** of astrocytes transfected with the indicated siRNA and migrating in a scratch wound assay. **(D)** Immunofluorescence images of wound-edge astrocytes transfected with the indicated siRNA and fixed 8h after wounding. Images show microtubules (anti-tubulin, white), *cis*-Golgi (anti-GM130, green), centrosome (anti-pericentrine, red) and the nucleus (DAPI, blue). **(E)** Quantification of Golgi orientation in astrocytes transfected with the indicated siRNA, 8h after wounding. The red line indicate the values expected for random positioning of the Golgi apparatus. **(F)** Fluorescence images showing PKCζ localization in wound-edge astrocytes transfected with the indicated siRNA. **(G)** Percentage of wound-edge cells showing PKCζ accumulation at the cell front. **(H)** Quantification of Golgi reorientation in a rescue experiment using astrocytes transfected with control siRNA or with a siRNA targeting both Cdc42 isoforms together with the indicated control (GFP) or GFP-tagged CDC42 constructs. Graphs show data presented as means ± SEM of 5 **(C)**, 4 **(E, G)** and 3 **(H)** independent experiments. Atleast 250 cells (A-C) and 150 cells (D-H) were analysed per condition. Ctl: Control cells transfected with non-relevant siRNA. p-values were calculated using two-sided unpaired Student’s t-test. Scale bars: 10 µm.

Associated with the alteration of directional persistence, the number of filopodia, classic actin-driven function of CDC42 at the cell leading edge was strongly inhibited by the simultaneous depletion of both isoforms. A similar result was observed upon specific depletion of CDC42u. In contrast, CDC42b depletion alone had no impact on filopodia formation, suggesting a more predominant role of CDC42u in this phenomenon (Fig. S1E). To assess the mechanisms involved in the control of directed migration, we looked at Golgi reorientation in cells at the wound edge. Following wounding of the astrocyte monolayer, the Golgi apparatus together with the centrosome localizes in front of the nucleus in the direction of migration in a CDC42-dependent manner (Etienne-Manneville and Hall, 2001). Here, microtubule polarized rearrangement and Golgi reorientation were dramatically impaired by the depletion of both CDC42 isoforms as expected, and were similarly affected by the specific depletion of the CDC42u (Fig. 1D, E). In contrast, CDC42b knockdown had no detectable effect (Fig. 1D, E). The Par6/PKCζ polarity complex is a key mediator of CDC42 function in astrocyte polarization (Etienne-Manneville and Hall, 2001; Etienne-Manneville and Hall, 2003a; Etienne-Manneville and Hall, 2003b; Etienne-Manneville et al., 2005). Depletion of CDC42u alone or of both CDC42 variants, but not of CDC42b alone, prevented the recruitment of the aPKC protein family member PKCζ to the cell leading edge (Fig. 1F, G), confirming that CDC42u, but not CDC42b, is required for the activation of the polarity pathway in astrocytes. To determine whether the lack of function of CDC42b was due to its relatively low level of expression, siRNA-resistant CDC42u (u^RES^) or CDC42b (b^RES^) were expressed. u^RES^, but not b^RES^ rescued Golgi reorientation in CDC42-depleted astrocytes (si-both) (Fig. 1H). Together, these observations point to the ubiquitous CDC42u isoform as the main regulator of cell polarity and directed migration in astrocytes.

### Brain and ubiquitous isoforms show similar binding partners

We then investigated the molecular differences that could explain the functional specificity of the two CDC42 isoforms. The alternative splicing of the carboxy-terminal (C-ter) exons causes a major difference in the last 10 amino acids of CDC42 isoforms (Fig. 2A). Since the C-ter domain of Rac was shown to participate in its interaction with some of its effectors (Abdrabou and Wang, 2018; Knaus et al., 1998), we examined whether both CDC42 variants could interact with the same effectors. We first tested their interaction with Par6 and PKCζ by performing immunoprecipitation of constitutively active (CA) V12-, GFP-tagged CDC42b and CDC42u in HEK cells (Fig. 2B). In these conditions, both isoforms appear to interact similarly with the Par6-aPKCζ polarity complex (Fig. 2C).

**Figure 2:**
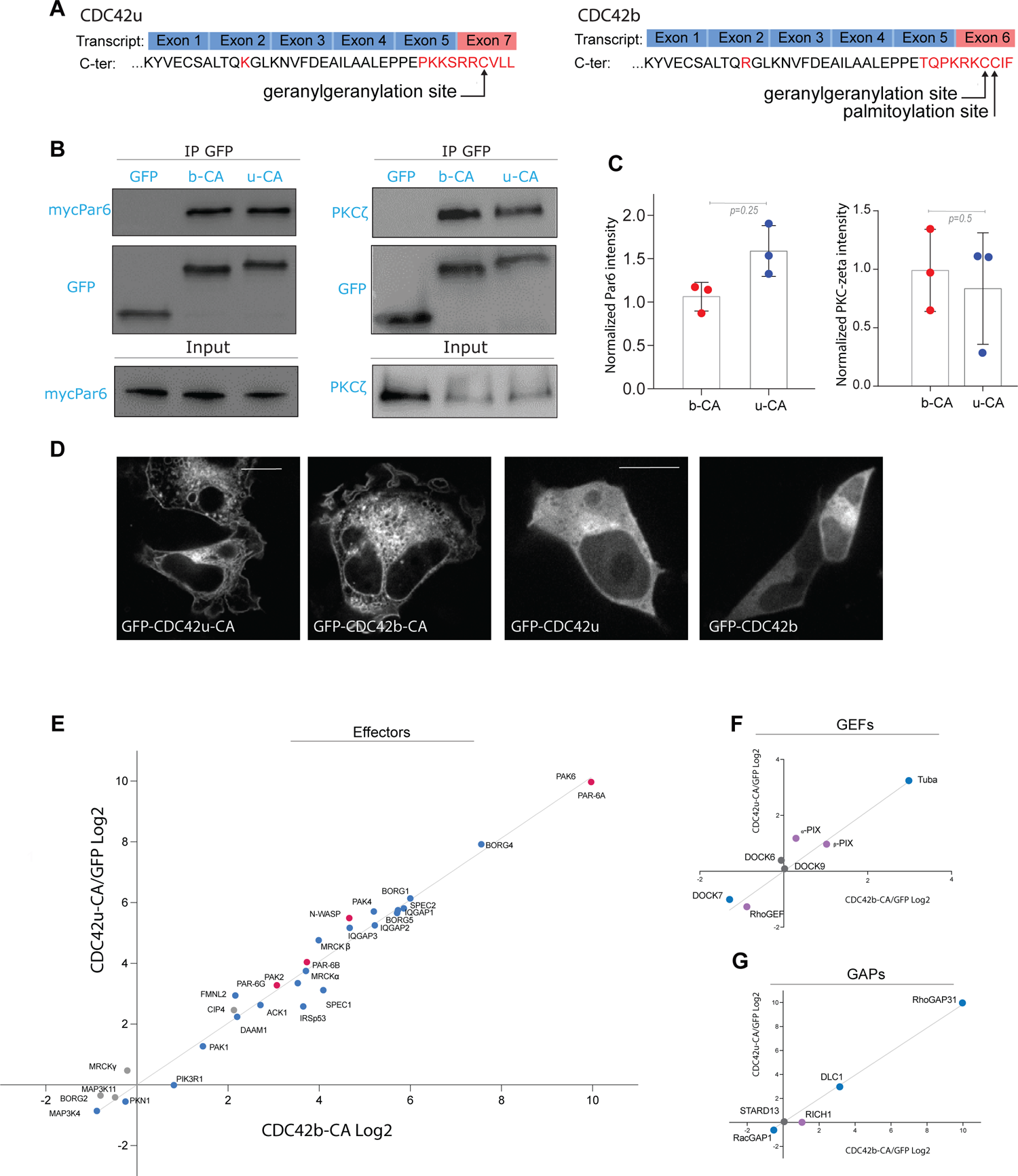
CDC42u and CDC42b share the same panel of binding partners. (A) Transcribed exons and translated carboxy-terminal protein sequences of ubiquitous (CDC42u) and brain CDC42 (CDC42b). (B) Western blots showing immunoprecipitation of GFP-tagged proteins; GFP-CDC42b CA (b-CA) and GFP-CDC42u CA (u-CA) from transfected HEK cells. Co-immunoprecipitation of mycPar6 and PKCζ are shown in the top panels and their respective expression levels in total cell lysate in the lower input panel. (C) Quantification of co-immunoprecipitated mycPar6 and PKCζ normalized to their input values. Graph shows data points and means ± SEM of 3 independent experiments. (D-F) Correlation analysis plots showing the fold change (in Log2 units) in number of peptides of a given protein co-precipitated with constitutively active (CA) CDC42b (x-axis) versus with CDC42u (y-axis). The number of peptides is normalized by the respective number obtained in GFP control immunoprecipitation. Proteomic screening was performed by applying loose filtering parameters to segregate the interactors of the CA screen into GEFs, GAPs and effectors. We identified 29 effectors out of 40 known epithelial effector proteins of CDC42. N-WASP and PAR6 proteins have been highlighted in red (E), 7 known GEFs (F) and 5 known GAPs (G). (D-F) Peptides used to calculate significance are ≥ 6, p-value ≤ 0.05 and number of replicates = 4. (G) Spinning disk images of HEK cells overexpressing the indicated pEGFP-CDC42 constructs 24h after transfection. Note that, in these conditions, CDC42u and CDC42b diplay similar localization pattern

Thus, we next performed GFP-trap pull-down assays using HEK cells overexpressing GFP tagged CDC42 isoforms (Fig.2D). The GFP-tagged WT- or CA-Cdc42 resins along with co-immunoprecipitated interactors were sent for a mass spectrometric proteomic screen. Quantitative analysis was performed to assess the fold change of peptides bound to one isoform of CDC42 in comparison to GFP control. Analysis of best binding partners was performed with the CA mutant of each isoform to capture a maximum of effector proteins (see methods for access to PRIDE repository containing raw data and screen analysis). We then performed loose filtering parameters to segregate the interactors of the screen into effectors and also GEFs, GAPs. We identified 29 effectors out of 40 known epithelial effector proteins of CDC42 (Fig 2D, S2A) (Pichaud et al., 2019). We could also detect 7 known GEFs (Fig 2E, S2B) and 5 known GAPs (Fig 2F, S2C). We could not identify any statistically significant difference in the list of effectors or in the fold change of effectors between both isoforms (Fig. S2 and PRIDE repository). These results show that despite their divergent carboxy-terminal sequences, both CDC42 isoforms can biochemically interact with the same binding partners. From the correlation plots of the corresponding fold change for each isoform we analyzed the linear regression for all three plots. The R^2^ = 0.893 for GEFs, 0,900 for GAPs and 0.978 for effectors therefore indicated a good linear fit correlation (Fig 2D-F), which may suggest that the alternative splicing does not strongly modify the binding affinity of the CDC42 with its main partners.

### Brain and ubiquitous CDC42 display different intracellular localization

Seeking an alternative explanation to the functional specificity of the two isoforms, we asked whether the carboxy-terminal sequence which includes the CAAX box required for membrane recruitment of the CDC42 proteins may influence the protein subcellular localization and ability to encounter its interactors. Immunoprecipitation experiments are typically performed using cells which strongly overexpress the tagged CDC42 (Fig. 2G), leaving the possibility that a specific subcellular localization of endogenous protein isoforms may be a critical factor in the control of CDC42 interactions with its effectors.

Since we could not find nor produce any antibody that specifically recognize CDC42 isoforms in immunoprecipitation or in immunofluorescence, we microinjected constructs encoding fluorescently-tagged CDC42 isoforms into astrocyte nuclei. Cells were imaged after a short time of expression (4h) to obtain low levels of expression. In contrast to cell transfection, short-term expression after microinjection revealed a different localization of the two CDC42 isoforms in astrocytes (Fig. 3A and Movie 1-3). Movie 1 shows non-migrating astrocytes expressing GFP-CDC42b together with mCherry-CDC42u. CDC42u is mainly cytosolic, whereas CDC42b appears to be largely associated with intracellular compartments. Movies 2 and 3 show the localization of CDC42 isoforms in cells actively migrating in a scratch-induced migration assay, previously shown to activate CDC42 through the integrin-mediated recruitment of the GEF βPIX (Osmani et al., 2010).. CDC42u accumulated at the leading edge plasma membrane, including in filopodia. CDC42b was recruited to the cell front, generally slightly behind the leading edge (Fig. 3A). CDC42b was most strikingly found on intracellular membranes including cytoplasmic vesicles where it colocalized with the early endosome marker EEA1 and Golgi apparatus where it colocalized with the cis-marker GM130 (Fig. 3A-C). In contrast, CDC42u was rarely detectable at these sites (Movie 3, Fig. 3A-C). This differential subcellular localization of the isoforms was confirmed using a different, smaller ALFA tag (Fig. S1F). We asked whether specific subcellular localization was due to the divergent C-ter domain of CDC42. The C-ter CAAX box of both isoforms is geranyl-geranylated but CDC42b can bear an additional palmitoyl group attached to the last cysteine residue in a reversible manner (Kang et al., 2008; Nishimura and Linder, 2013, Nishimura, 2019 #19536). Suppression of the geranyl-geranylation by a CVLL to SVLL mutation in CDC42u (u(SVLL)) (Nishimura and Linder, 2013) led to its accumulation in the nucleus (Fig. 3D). A similar CCIF to SCIF mutation in CDC42b (b(SCIF)) to prevent all lipid modification of CDC42b resulted in complete loss of endomembrane-binding and cytosolic distribution of the protein. In contrast, the CCIF to CSIF mutation, which specifically prevents palmitoylation but not geranyl-geranylation, inhibited recruitment of CDC42b to the plasma membrane and on endocytic vesicles, but did not prevent its association with the Golgi apparatus (Fig.3D), suggesting that palmitoylation controls the recruitment of CDC42b to the cytoplasmic vesicles from the Golgi apparatus. We confirmed the role of lipid anchors in the membrane association of CDC42 isoforms by treating cells with GGTI298, which prevents both geranyl-geranylation and palmitoylation, or with 2-bromopalmitate (2BP), a specific inhibitor of palmitoylation (Fig. S3A). These results show that the lipid modifications of the carboxy-terminal domain of CDC42 isoforms are crucial for the specific membrane association of the two proteins. They further suggest that the divergent C-ter domains may contribute to the functional specificities of CDC42 isoforms by controlling their subcellular localization, indirectly affecting their abitlity to interact with their effectors.

**Figure 3:**
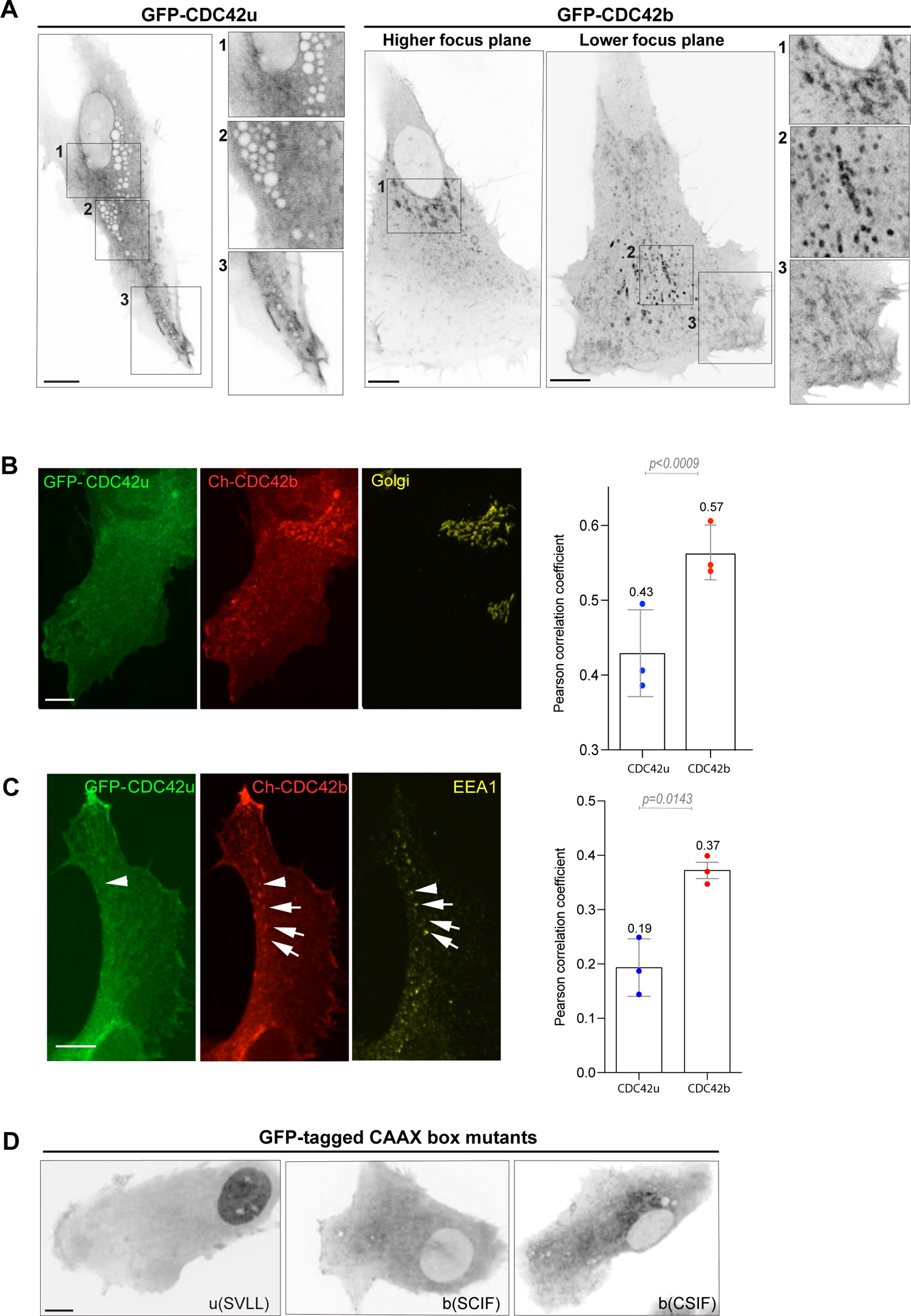
Localization of brain CDC42 (CDC42b) and ubiquitous CDC42 (CDC42u) in migrating astrocytes. **(A)** Confocal section images of migrating astrocytes 3 to 4h following microinjection of GFP-tagged CDC42u or CDC42b constructs and 5h after wounding. Right panels show higher magnification of the corresponding boxed area and highlight the different localization of the two isoforms (see corresponding movies 2 and 3). **(B)** Confocal stack images of GFP-CDC42u and mCherry-CDC42b expressing astrocytes 5h after wounding stained with anti-GM130 (cis-Golgi marker). Right panel, quantification of the colocalisation of each CDC42 construct with GM130. **(C)** Confocal stack images of GFP-CDC42u and mCherry-CDC42b expressing astrocytes 5h after wounding stained with anti-EEA1 (early endosomes marker). Right panel, quantification of the colocalisation of each CDC42 construct with EEA1. **(D)** Confocal section images showing localization of GFP-tagged non-lipid-modified ubiquitous (u(SVLL)) or brain CDC42 (b(SCIF)) and of non-palmitoylatable CDC42b (bCSIF) in migrating astrocytes 5h after wounding. All graphs show the values and means ±SEM of 3 independent experiments and atleast 30 cells were analysed per condition. All p-values were calculated using two-sided unpaired Student’s t-test. Scale bars: 10 µm.

### Brain CDC42 is the major CDC42 isoform involved in N-WASP dependent endocytosis

Following this line of reasoning, we sought for CDC42b specific functions on intracellular membrane compartments. In migrating astrocytes, CDC42 not only controls Par6-dependent cell polarity, it is also involved together with Arf6 in vesicular recycling from the plasma membrane (Osmani et al., 2010). Because CDC42b showed a strong enrichment on EEA1-positive organelles compared to CDC42u (Fig. 3A-C), we performed dextran uptake experiments to examine the role of CDC42 isoforms in the formation of the EEA1-positive pinosomes which are frequently observed at the leading edge of migrating astrocytes. Knockdown of both isoforms of CDC42 strongly reduced (−47%) dextran uptake (Fig. 4A). Strikingly, CDC42u depletion caused a minor reduction of dextran internalization, whereas CDC42b knockdown significantly decreased the uptake rates (by approximately 40%; Fig. 4A). The predominant role of CDC42b in pinocytosis was confirmed by rescue experiments in astrocytes depleted for both isoforms. GFP-CDC42b^RES^ led to a stronger rescue than GFP-CDC42u^RES^ (Fig. 4B). The non-lipid-modified mutants of either isoform (b^RES^(SCIF), u^RES^(SVLL)) did not rescue dextran uptake (Fig. 4B). Furthermore, overexpression of the non-palmitoylable CDC42b mutant (b^RES^(CSIF)) did not restore the dextran uptake (Fig. 4B) indicating that palmitoylation, which promotes CDC42b association with intracellular vesicles specifically (Fig. 3D) is crucial for its function in pinocytosis. Dextran uptake experiments using specific inhibitors of lipid modification (GGTI298 or 2BP) confirmed these findings (Fig. S3B). These results show that palmitoylated CDC42b is the major CDC42 isoform involved in pinocytosis in migrating astrocytes.

**Figure 4:**
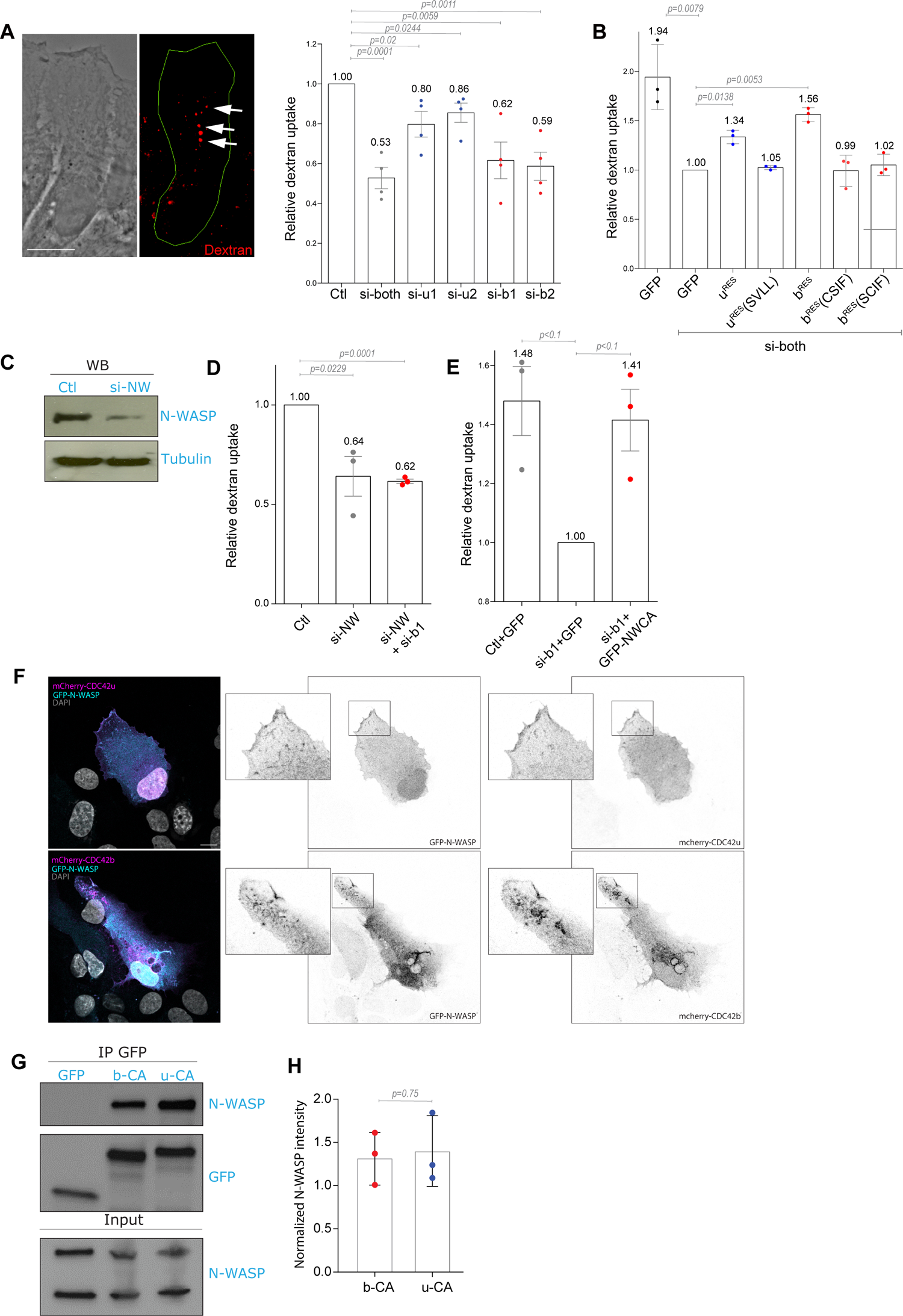
CDC42b, N-WASP, but not CDC42u control pinocytosis. **(A)** Phase and fluorescence images showing dextran uptake in control migrating astrocytes. Right panel, Quantification of the relative dextran uptake in astrocytes nucleofected with the indicated CDC42 siRNA. **(B)** Quantification of the relative dextran uptake in a rescue experiment using control or CDC42-depleted astrocytes expressing the indicated CDC42 construct. **(C)** Western Blot analysis of N-WASP expression in control (Ctl) and N-WASP siRNA nucleofected astrocytes. Tubulin was used as loading control. **(D)** Quantification of the relative dextran uptake in astrocytes nucleofected with indicated siRNAs (NW: N-WASP). **(E)** Quantification of relative dextran uptake in a rescue experiment using control or CDC42b-depleted astrocytes expressing GFP or GFP-N-WCA (constitutively active N-WASP) constructs. All graphs show the values and means ±SEM of 3 independent experiments with at least 150 cells analyzed per condition. Data were normalized to the values obtained for si-both treated and GFP expressing cells. **(F)** Confocal section images from migrating astrocytes showing co-localization of mCherry-CDC42b and GFP-N-WASP at the wound edge of migrating astrocytes and specifically with vesicles 6h after wounding while Cdc42u colocalized with N-WASP puncta, macropinosomes were absent. **(G)** Western blot showing immunoprecipitation of GFP-tagged proteins; GFP-CDC42b CA (b-CA) and GFP-CDC42u CA (u-CA) from transfected HEK cells and co-immunoprecipitation of N-WASP. **(H)** Quantifications of co-immunoprecipitated N-WASP normalized to its respective expression levels in total cell lysate (input). All p-values were calculated using two-sided unpaired Student’s t-test. Scale bars: 10 µm.

SiRNA-mediated depletion of N-WASP (Fig. 4C) inhibited dextran uptake in migrating astrocytes to a similar level as CDC42b knock-down (Fig. 4A and 4B), as previously reported (Kessels and Qualmann, 2002; Legg et al., 2007). A double knockdown of CDC42b and N-WASP did not further increase the reduction in dextran uptake (∼40%) (Fig. 4D). Finally, expression of an activated form of N-WASP (GFP-NWCA) in CDC42b depleted cells rescued macropinocytosis confirming that N-WASP is the major effector of CDC42b controlling macropinocytosis in astrocytes (Fig. 4E). We looked at the localization CDC42b colocalized with N-WASP on macropinosomes in migrating astrocytes, whereas CDC42u was rarely observed on any intracellular vesicles including macropinosomes (Fig. 3A, 3C and 4F, movie 4). These data indicate that the C-ter domain of CDC42b controls its association with intracellular vesicles, its interaction with N-WASP and CDC42/N-WASP-mediated pinocytosis. However, when N-WASP association with CDC42 isoforms was assessed in overexpressing HEK cells (Fig. 4G), there was no significant difference in the ability of N-WASP to bind each isoform (Fig. 4G, 4H) as observed in proteomic analysis of CDC42 interactome (Fig. 2E). We conclude that even if both isoforms can interact with N-WASP in vitro, CDC42b is main regulator of N-WASP-dependent micropinocytosis in cells because of its specific localization on endocytic vesicles.

### Both CDC42 isoforms contribute their specific functions during chemotaxis of neural precursor cells

Since CDC42 isoforms show functional differences, we asked whether they may cooperate in more complex migratory situations where both front-rear polarization and endocytosis are required. Neural precursor cells (NPCs) migrate long distances from their zones of origin to their final destination where they differentiate, following gradients of chemoattractants (Leong et al., 2011). In NPCs, both isoforms are expressed at approximately identical levels (Fig. 5A) (Yap et al., 2016). Like in astrocytes, short-term expression of microinjected CDC42 constructs in NPCs illustrated the distinct subcellular localization of CDC42u and CDC42b. CDC42u was mainly visible in the cytosol and at the plasma membrane and the CDC42b accumulated on intracellular EEA1-positive vesicles and on the Golgi apparatus (Fig. 5B, 5C).

**Figure 5:**
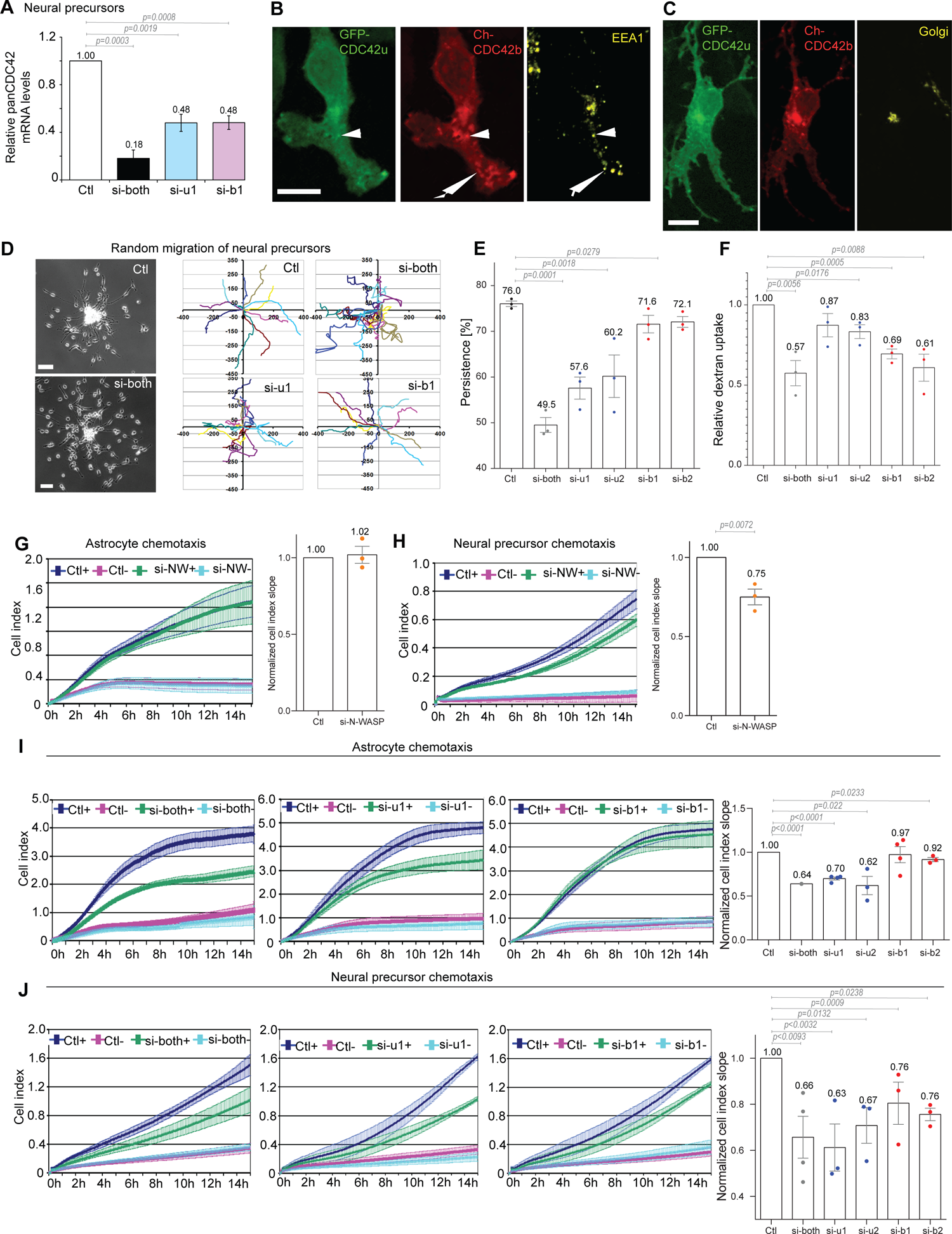
CDC42 isoforms cooperate to promote neural precursor cell (NPC) chemotaxis. **(A)** Total CDC42 mRNA levels were measured using a TaqMan assay that recognizes both CDC42 isoforms (panCDC42) in neural precursors transfected with the indicated siRNA. **(B, C)** Fluorescence images of NPCs expressing GFP-CDC42u and mCherry-CDC42b, fixed and stained with anti EEA1 (early endosome marker) **(B)** and anti-GM130 (Golgi) **(C)**. **(D)** Phase contrast images of NPCs nucleofected with the indicated siRNA and migrating out of neurospheres 5h after plating. Right panels show the representative cell trajectories over 4h of migration. **(E)** Directional persistence of NPC migration measured between 100min and 300min after plating. **(F)** Relative dextran uptake into NPCs transfected with the indicated siRNAs. The graph shows the data and the means ± SEM of 3 independent experiments with at least 150 cells analyzed per condition. **(G-J)** Astrocyte and NPC chemotaxis in Boyden chamber-based xCelligence system assays. +: bottom well contains FBS; -: no FBS in bottom well. Curves show the impedance measurement over time on the bottom surface of the filter. Right panels show the curve slopes, indicative of the rate of migration in chemotactic conditions in which FBS was contained in the bottom wells. Graphs show the data and the means ± SEM of at least 3 independent experiments. Data were normalized to the control. **(G-H)** Effects of N-WASP (si-NW) knockdown on chemotactic migration of astrocytes **(G)** or NPCs **(H)**. **(I-J)** Chemotactic migration of astrocytes **(I)** or NPCs **(J)** upon knockdown of CDC42 isoforms. All p-values were calculated using two-sided unpaired Student’s t-test. Scale bars: 100 µm.

NPCs grow in a primary 3D tissue culture system (neurospheres) and their migration can be observed when neurospheres are placed on an adhesive substrate (Durbec et al., 2008) (Fig. 5D). Using video microscopy, we first analyzed NPC migration out from neurospheres. The directional persistence was strongly reduced in CDC42u-depleted cells (from 76% to 58-60%) and in cells lacking both isoforms (to 50%), but was not significantly altered by CDC42b or N-WASP depletion (Fig 5D, 5E, and S3C).

We next performed dextran uptake assays in siRNA treated NPCs. CDC42b specific depletion, like N-WASP depletion, significantly decreased dextran uptake into NPCs, whereas knockdown of CDC42u had no significant effect (Fig. 5F and S3D). These results in NPCs confirmed the predominant role of CDC42u in cell polarization and of CDC42b in macropinocytosis.

We then tested the contribution of the two isoforms during NPC chemotactic migration, during which endocytosis is involved in the processing of chemotactic signals (Zhou et al., 2007). We used a Boyden chamber-based xCelligence-assay to analyze the chemotactic migration of astrocytes and NPCs. In the absence of a gradient, astrocytes and NPCs barely migrated through the filter (Fig. S3E, S3F). Addition of FBS in the lower compartment induced the chemotactic migration of both astrocytes and NPCs (Fig. 5G, 5H). The knockdown of N-WASP decreased chemotaxis efficiency in NPCs but did not affect astrocyte chemotaxis (Fig. 5G, 5H). When astrocytes or NPCs were transfected with siRNAs targeting both CDC42 isoforms, chemotaxis was strongly reduced in both cell types (Fig. 5I, 5J). For both cell types, CDC42u specific depletion also strongly reduced chemotactic migration. In contrast, CDC42b depletion led to a significant decrease in N-WASP-dependent NPC chemotaxis without altering N-WASP-independent astrocyte migration (Fig. 5I, 5J). We conclude that CDC42b is the major isoform involved in N-WASP dependent function in NPC chemotaxis and that both isoforms of CDC42 contribute their specific functions to participate to complex migratory processes such as NPC chemotaxis.

## Discussion

In conclusion, we show that, while both CDC42 isoform can in principle interact with the same effectors in vitro, they do have non-redundant functions in cells. These findings illustrate the impact of subcellular localization on specific protein molecular interactions and cellular functions. More specifically, efficient cell polarization and directed persistent migration solely require the CDC42u isoform both in astrocytes and in NPCs. Although both CDC42u and CDC42b can biochemically interact with Par6-aPKC, only CDC42u can recruit Par6 and aPKC at the leading edge plasma membrane. Conversaly, although both CDC42u and CDC42b can biochemically interact with N-WASP, CDC42b which preferentially localizes on intracellular compartments, is the predominant splice variant involved in the regulation of N-WASP-dependent endocytosis. Altogether, these observations the importance of subcellular compartmentalization in CDC42 and more generally Rho GTPases’ functions. The recruitment of CDC42 as well as other RhoGTPases is dictated by their carboxy-terminal sequence and at least in part by lipid modifications (Farhan and Hsu 2016; Ravichandran, Goud, and Manneville 2020). Related to this finding, mutations that touch on this carboxy-terminal variable sequence of CDC42 and can affect either one or both CDC42, are associated with distinct dramatic disorders (Bekhouche et al., 2020; Martinelli et al., 2018).

The fact that palmitoylation as well as scratch-induced signaling influence Cdc42b recruitment to trafficking vesicles from the Golgi apparatus suggests that each isoform may be regulated by distinct pathways. The fact that GDI3, a GDI expressed in the brain and pancreas, localizes at the Golgi apparatus via a N-terminal hydrophobic anchor (Brunet et al., 2002), suggests yet another potential regulatory mechanism. Future work will focus on deciphering how the amino acid sequence and/or the lipid modifications lead to such radical differences in CDC42 localization to better understand how alterations of the carboxy-terminal sequence can induce specific diseases (Bekhouche et al., 2020).

The two CDC42 isoforms have been reported to be functionally specialized during neurogenesis. CDC42u stimulates mTORC1 activity to promote neuroprogenitor formation, whereas CDC42b works synergistically with activated CDC42-associated kinase (ACK) in down-regulating mTOR expression and promoting neuronal differentiation. It is remarkable that the two highly-similar CDC42 splice variants regulate distinct stages of neurogenesis and neuronal differentiation (Endo et al., 2020). Moreover the conditional inactivation of the *CDC42* gene in cortical neurons reduces the efficiency of axon formation (Garvalov et al., 2007). In this context, CDC42u whose mRNA preferentially localizes into axons plays a role in axonogenesis (Lee et al., 2021) whereas the palmitoylation of CDC42b accounts for its preferential localization to dendritic spines and its role in dendrite maturation. These findings have advanced the understanding of mechanisms underlying axo-dendritic polarity in developing neurons and argue that co-expression of the non-redundant CDC42 isoforms in the same cell is important during neuronal development (Yap et al., 2016)..With the identification of the role of CDC42b in the formation of dendritic spines during neuronal differentiation (Kang et al., 2008; Wirth et al., 2013), our identification of its role in endocytosis and NPC chemotaxis is increasing evidence that the brain variant controls specific neural functions. NPC migration is a crucial step in brain development and conditional deletion of both CDC42 isoforms in mouse NPCs has been shown to cause malformations in the brain (Chen et al., 2006). Nevertheless both CDC42 isoforms contribute, albeit differently, to the behavior of NPCs, underlining that the co-expression of these non-redundant isoforms is essential during neuronal development.

## Materials and Methods

### Antibodies and inhibitors

The following primary antibodies were used in this study: rat monoclonal anti-α-tubulin (AbDSerotec MCA77G), mouse monoclonal anti-GM130 (BD Transduction 610823), rabbit polyclonal anti-pericentrin (Covance PRB 432-C), rabbit polyclonal anti-N-WASP (Abcam AB23394), rabbit polyclonal anti-PKCζ C-20 (Santa Cruz Biotech SC-216), mouse anti-EEA1(BD biosciences 610457) and HRP coupled anti-GFP (Abcam ab6663). As secondary antibodies we used standard antibodies from Jackson ImmunoResearch: Cy5 conjugated donkey anti-mouse, Alexa Fluor 488 conjugated donkey anti-rat, TRITC conjugated donkey anti-rabbit, as well as HRP coupled donkey anti-mouse, anti-rabbit and anti-goat. For Western Blot detection of CDC42b a custom-made isoform specific rabbit polyclonal antibody was generated by Covalab using 2 peptides for 2 immunization steps. Peptide 1 for first immunization step: C-AALEPPETQPKRK-coNH2 (CDC42b amino acids 175-187); peptide 2 for second immunization step: C-ETQPKRK-coNH2 (CDC42b amino acids 181-187). The antibody was subsequently purified from the anti-serum via immobilized peptide 2. DAPI in ProLong Gold Antifade Reagent (Life Tech) was used to visualize nuclei. To suppress lipid modification of CDC42 isoforms, cells were treated overnight in 120 µM 2BP (to suppress palmitoylation; Sigma) or 20 µM GGTI298 (to suppress geranyl-geranylation and palmitoylation; Tocris)

### Cell Culture

All procedures were performed in accordance with the guidelines approved by the French Ministry of Agriculture, following European standards. Preparation of neurosphere cultures was performed as described (Calaora et al., 2001). Briefly, the striata of E14 OFA rats were removed from the embryos and mechanically dissociated before cells were seeded at 1.2 × 10^5^ cells/ml in uncoated 260 ml culture flasks (Fisher Bioblock Sc.). Culture medium consisted of DMEM/F-12 (Gibco) supplemented with 2% B27 (Gibco) and 50µg/ml gentamicin (Sigma) in the presence of 20 ng/ml EGF (R&D Systems Europe). Media were supplemented with 20 ng/ml EGF every 48h, and spheres passaged using 0.025% trypsin-EDTA (Gibco) on the fourth and sixth day in culture. Human FGF-b (RayBiotech) was also added to the medium at 10 ng/ml for the first four days of culture. For preparation of primary astrocyte cultures, the telencephala of E18 OFA rats were removed from the embryos and mechanically dissociated. Cells were plated and maintained as previously described (Etienne-Manneville, 2006) using 1g/l glucose DMEM (Gibco) supplemented with 10% FBS (Eurobio) and penicillin/streptomycin (10,000 U ml^−1^ and 10,000 μg ml^−1^; Gibco) as culture medium. HEK and HeLa cells were cultured in 4.5g/l glucose DMEM (Gibco) supplemented with FBS and antibiotics as added for astrocytes. All cells were kept in a 37°C incubator at 5% CO2.

### Immunoprecipitation assay and mass spectrometric proteomic screen

GFP Immunoprecipitation assay was carried out, where HEK cells transfected with GFP tagged constructs of CDC42 were lysed using 50mM TRIS base, Triton 2 %, 200mM NaCl as well as 1 tablet/10 ml protease inhibitor Mini-complete, EDTA-free (Roche). After removal of insoluble fragments via centrifugation at 12,000 g for 25 min, lysates were incubated with 15 µl of GFP-Trap Agarose beads from Chromotek for 1h at 4 °C on a rotary wheel. The beads were washed using a wash buffer comprising of 50mM TRIS base, 150mM NaCl, 1mM EDTA and 2.5 mM MgCl2 and pH adjusted to 7.5. Following the final wash beads were stored with wash buffer in 4°C prior to depositing at the Institut Curie Mass Spectrometry and Proteomics facility (LSMP).

Where proteins on beads were washed twice with 100μL of 25mM NH_4_HCO_3_ and we performed on-beads digestion with 0.2μg of trypsin/LysC (Promega) for 1 hour in 100µL of 25mM NH_4_HCO_3_. Sample was then loaded onto a homemade C18 StageTips for desalting. Peptides were eluted using 40/60 MeCN/H2O + 0.1% formic acid and vacuum concentrated to dryness.

Online chromatography was performed with an RSLCnano system (Ultimate 3000, Thermo Scientific) coupled to an Orbitrap Fusion Tribrid mass spectrometer (Thermo Scientific). Peptides were trapped on a C18 column (75μm inner diameter × 2cm; nanoViper Acclaim PepMapTM 100, Thermo Scientific) with buffer A (2/98 MeCN/H_2_O in 0.1% formic acid) at a flow rate of 4.0µL/min over 4 min. Separation was performed on a 50cm x 75μm C18 column (nanoViper Acclaim PepMapTM RSLC, 2μm, 100Å, Thermo Scientific) regulated to a temperature of 55°C with a linear gradient of 5% to 25% buffer B (100% MeCN in 0.1% formic acid) at a flow rate of 300nL/min over 100 min. Full-scan MS was acquired in the Orbitrap analyzer with a resolution set to 120,000 and ions from each full scan were HCD fragmented and analyzed in the linear ion trap.

For identification the data were searched against the Homo sapiens (UP000005640) SwissProt database using Sequest HF through proteome discoverer (version 2.2). Enzyme specificity was set to trypsin and a maximum of two missed cleavage site were allowed. Oxidized methionine, N-terminal acetylation, and carbamidomethyl cysteine were set as variable modifications. Maximum allowed mass deviation was set to 10 ppm for monoisotopic precursor ions and 0.6 Da for MS/MS peaks.

The resulting files were further processed using myProMS (Poullet et al., 2007) v3.6 (work in progress). FDR calculation used Percolator and was set to 1% at the peptide level for the whole study. The label free quantification was performed by peptide Extracted Ion Chromatograms (XICs) computed with MassChroQ version 2.2 (Valot et al., 2011). For protein quantification, XICs from proteotypic peptides shared between compared conditions (TopN matching) with no missed cleavages were used. Median and scale normalization was applied on the total signal to correct the XICs for each biological replicate. To estimate the significance of the change in protein abundance, a linear model (adjusted on peptides and biological replicates) was performed and p-values were adjusted with a Benjamini–Hochberg FDR procedure with a control threshold set to 0.05. The mass spectrometry proteomics data have been deposited to the ProteomeXchange Consortium via the PRIDE (Vizcaino et al., 2016) partner repository with the dataset identifier PXD017477(username, Password: 4fNr03LX)

### Cell transfection and RNAi

siRNA constructs were introduced into rat astrocytes or NPCs by Nucleofection technology (Amaxa Biosystems) using Lonza protocols. Plasmids encoding fluorescently tagged constructs were microinjected. All siRNAs were obtained from Eurofins except for a non-targeting control, which was obtained from Dharmacon. To quantify knockdowns, protein samples were analyzed using ECL immunoblotting and ImageJ. Alternatively, mRNA levels were measured using qPCR (see below). In each case, cells were analyzed 4 days after transfection with siRNAs. Transfection of HEK293 and HeLa cells with plasmids was performed with the calcium phosphate method.

si-N-WASP: 5’-CUUGUCAAGUAGCUCUUAA(dTdT)-3’

si-both: 5’-UGAUGGUGCUGUUGGUAAA(dTdT)-3’

si-u1: 5’-CAAUAAUGACAGACGACCU(dTdT)-3’

si-u2: 5’-GCAAUAUUGGCUGCCUUGGUU(dTdT)-3’

si-b1: 5’-CCAUUUAACAAUCGACUUA(dTdT)-3’

si-b2: 5’-ACUCAACCCAAAAGGAAGUUU(dTdT)-3’

### Real time qPCR

For isolation of total RNA striatal neurospheres or cultured astrocytes were prepared from E14 or E18 rat embryos respectively. RNA was isolated by using the RNeasy Mini Kit (QIAGEN) followed by the digestion of contaminating genomic DNA (Turbo DNA Free). RNA concentration and purity were determined by spectrophotometry. cDNA synthesis was performed according to kit instructions (VILO cDNA synthesis; Invitrogen). Quantitative real-time PCR was performed using Applied Biosystems custom TaqMan Gene Expression Assays designed using the Taqman Assay Search Tool (Life Technologies) for the ubiquitous CDC42 isoform. Assay Rn00821429_g1 was used to quantify panCDC42 mRNA levels (both isoforms). As endogenous control we used PGK1 (assay Rn00821429_g1) and Ppia (assay Rn00690933_m1) (all from Life Technologies) for astrocytes and Casc3 (Rn00595941_m1) and Eif2b1 (Rn00596951_m1) for neural precursor cells. Real-time PCR amplification was performed on a Sequence Detection System (7500; Applied Biosystems) using TaqMan Universal MMix II (Life Technologies) according to the manufacturer’s instructions. Thermal cycling conditions were as follows: 50°C for 2 min, 95°C for 10 min followed by 40 cycles of 95°C for 15 s, and 60°C for 1 min. Data were collected and analyzed with the SDS software v2.0.6 (Applied Biosystems). The comparative CT Method (ΔΔC_T_) was used as described in the AB7500 SDS guidelines.

### Live spinning disk confocal microscopy

To study protein localization in live cells, plasmids encoding the GFP- or mCherry tagged protein were microinjected into wound border astrocytes plated on glass bottom dishes (MatTek) and scratched 1h before microinjection. Live imaging was done 4-8h later using a spinning disk confocal microscope (Perkin Elmer Ultra View ERS) equipped with a heating chamber (37°C) and CO2 supply (5%).

### Centrosome/Golgi reorientation and PKCζ localization assays

Scratch-induced cell polarization of astrocytes was studied as detailed previously (Etienne-Manneville, 2006). Briefly, a monolayer of astrocytes plated on poly-L-ornithine (Sigma) coated coverslips was scratched and fixed 8h later followed by immunostaining of centrosome, Golgi and microtubules. Wound border cells were counted as polarized when the centrosome and Golgi were situated within the 90° or 120°C section of a virtual circle drawn around the nucleus that faces the wound (see Fig. 4B). Images were acquired on a Leica DM6000 epifluorescence microscope equipped with 40×, NA 1.25 and a 63×, NA 1.4 objective lenses and were recorded with a CCD camera (CoolSNAP HQ, Roper Scientific) using Leica LAS AF software. PKCζ accumulation at the cell front was studied in knockdown cells situated on poly-L-ornithine coated cover slips and fixed 4h after wounding.

#### Cell Spreading Assays

Sixteen-well E-plates (Ozyme) were pre-coated with 25µg/ml Fibronectin (Sigma) for NPCs or used uncoated for astrocytes. Single cell suspensions of 1×10^5^ NPC or 5×10^4^ astrocytes in 100µl of normal culture medium were added per well. Each condition (control or siRNA Transfected) was run in triplicate wells. Cell adhesion and spreading was measured as changes in impedance measured on the xCelligence RTCA DP Analyzer (ACEA Biosciences) according to the manufacturer’s instructions.

#### Cell Migration Assays

Astrocyte wound healing and neurosphere migration assays: For live imaging of astrocyte wound healing experiments, Transfected astrocytes were seeded into poly-L-ornithin (Sigma) coated 12-well standard plastic dishes using normal cell culture medium. Cell monolayers were scratched immediately before image acquisition followed by addition of 20mM Hepes (Sigma) as well as antioxidant (Sigma) to the medium and addition of liquid paraffin on top of the medium. Video time-lapse data were acquired on a Zeiss Axiovert 200M microscope equipped with a 37°C humidified heating chamber as well as CO2 supply (5%) by taking pictures every 15min over 16h and analyzed by manual tracking of cells using ImageJ, as described previously (Camand et al., 2012). For tracking of NPCs, neurospheres obtained from suspension cultures of transfected cells were plated into fibronectin coated 12-well plastic dishes immediately before life imaging followed by addition of HEPES, antioxidant and paraffin as for astrocytes. Images were taken every 5 min for NPCs.

#### Chemotaxis assays

The bottom wells of sixteen-well C.I.M plates (Ozyme) were filled with 160µl of normal culture medium for astrocytes or NPCs containing 10% FBS in both cases. The upper wells were filled with 50µl of normal culture medium without FBS (but containing B27 in case of NPCs). After 1h of equilibration in the incubator, single cell suspensions of 1×10^5^ NPC or 5×10^4^ astrocytes in 100µl medium (without FBS) were added to the upper wells plated in the upper wells. Each condition (control or siRNA Transfected) was run in triplicate wells on the xCelligence RTCA DP Analyzer according to user guidelines. Cell migration was measured as changes in impedance with the slopes of cell migration being compared over 10h.

#### Endocytosis assays

Neurospheres were incubated for 1.5h at 37°C in 1mg/ml Texas Red labeled dextran (10 kDa Invitrogen), extensively washed and plated on poly-L-ornithine and fibronectin coated glass coverslips for 4h, before being fixed in 4% paraformaldehyde. Transfected astrocytes were plated on poly-L-ornithine coated coverslips and scratched 4d later followed by addition of 1mg/ml fluorescent dextran. Cells were fixed in PFA after 1.5 h and analyzed on the Leica DM6000 epifluorescence microscope described above. The relative amount of dextran uptake was determined using ImageJ to quantify the number of fluorescence maxima per cell after background subtraction.

#### Pull down assays

HEK cells were used non-transfected to test interactions with an endogenous protein. Cells were lysed using pull down buffer containing 500mM NaCl, 15mM KCl, 8mM TRIS base, 12mM HEPES free base, 3mM MgCl2, 10% (v/v) glycerol, 1% (v/v) Triton X-100 as well as 1 tablet/10ml protease inhibitor Mini-complete, EDTA-free (Roche). After removal of insoluble compounds via centrifugation, lysates were incubated with glutathione Sepharose beads (GE Healthcare) coated with the other interaction partner for 30min at 4°C on a rotary wheel. Subsequently, beads were washed three times for 10min with the same buffer containing however doubled amounts of NaCl on the rotary wheel, followed by detection of associated proteins using Western Blotting.

## Supporting information

Figure S1rev

Figure S2rev

Figure S3rev

## Acknowledgements

This work was supported by the Centre National de la Recherche Scientifique and the Institut Pasteur. Y. Ravichandran was funded by the Polarnet ITN (Innovative Training Network) part of the European Commission and the Fondation pour la Recherche Médicale. J. Hänisch was funded by a Marie Curie post-doctoral grant. B. Boëda is a full-time INSERM researcher. We would like to thank members of the SEM lab for support and discussion. We gratefully acknowledge the Imagopole of Institut Pasteur (Paris, France).

The authors declare no conflict of interest.

## Abbreviations

CDC42u: Cdc42 placental isoform (ubiquitous)

CDC42b: Cdc42 brain isoform

u(SVLL): CAAX box mutant of CDC42u

b(SCIF), b(CSIF): CAAX box mutants of CDC42b

si-u1/si-u2: CDC42u specific siRNA

si-b1/si-b2: CDC42b specific siRNA

u^RES^: siRNA resistant CDC42u

b^RES^: siRNA resistant CDC42u

