## Supplementary figures and images for "The functional specificity of CDC42 isoforms is caused by their distinct subcellular localization"

### Figure S1rev

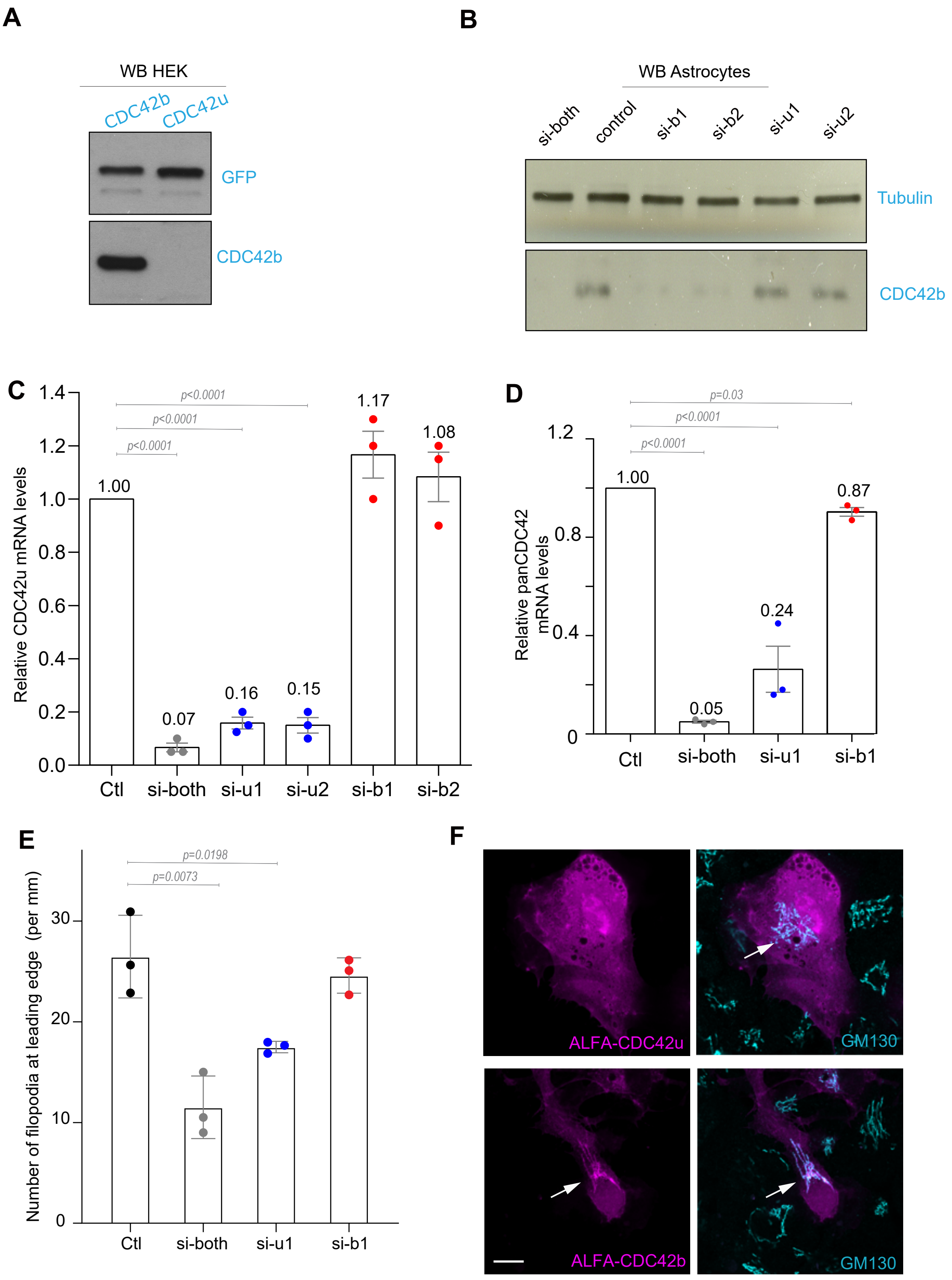


Supplementary Figure 1
Ravichandran et al.

### Figure S3rev

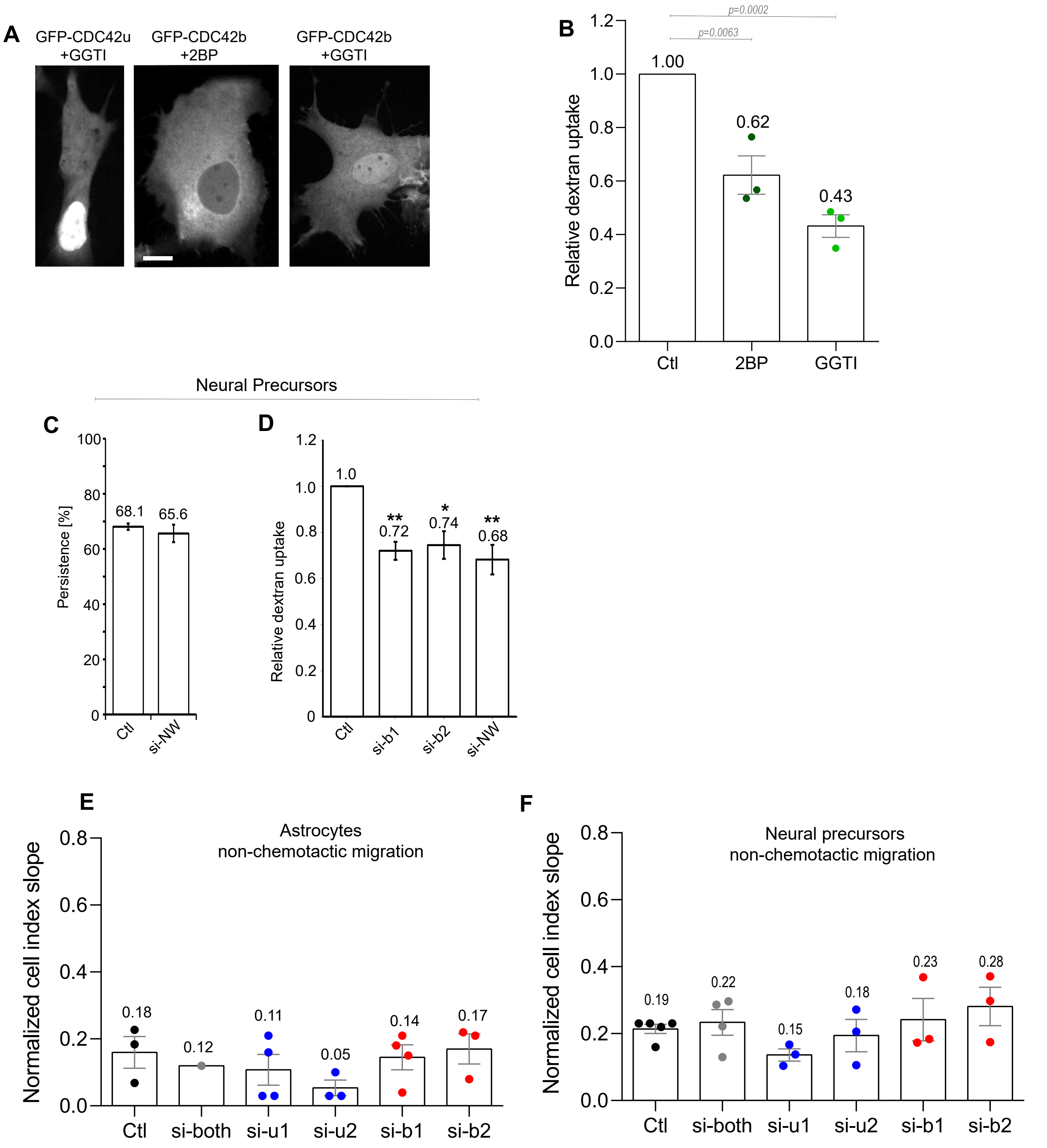


Supplementary Figure 3
Ravichandran et al.
