## Supplementary material for "The functional specificity of CDC42 isoforms is caused by their distinct subcellular localization": Figure S2rev

A

| Effectors | | | | | | | | |
|---|---|---|---|---|---|---|---|---|
| | Cdc42b-CA/GFP | | | | Cdc42u-CA/GFP | | | |
| Proteins | Ratio | Log2 | p-value | Peptides used | Ratio | Log2 | p-value | Peptides used |
| MAP3K4 | 0.54 | -0.88 | 1.27E-03 | 36 | 0.55 | -0.86 | 1.01E-04 | 31 |
| MRCKγ | 0.87 | -0.21 | 7.99E-01 | 29 | 1.39 | 0.47 | 5.54E-01 | 33 |
| IQGAP1 | 53.00 | 5.73 | 1.60E-188 | 430 | 53.74 | 5.75 | 3.14E-205 | 409 |
| MAP3K11 | 0.72 | -0.48 | 2.26E-01 | 8 | 0.75 | -0.41 | 2.44E-01 | 8 |
| PAR-6A | 1000.00 | 9.97 | | 27 | 1000.00 | 9.97 | | 26 |
| PAK6 | 1000.00 | 9.97 | | 8 | 1000.00 | 9.97 | | 6 |
| PKN1 | 0.84 | -0.25 | 2.56E-02 | 52 | 0.68 | -0.55 | 1.16E-04 | 55 |
| IQGAP2 | 37.33 | 5.22 | 1.39E-59 | 180 | 38.04 | 5.25 | 1.81E-68 | 169 |
| IQGAP3 | 25.53 | 4.67 | 8.25E-53 | 178 | 35.96 | 5.17 | 4.94E-59 | 170 |
| SPEC2 | 58.00 | 5.86 | 3.71E-08 | 16 | 55.95 | 5.81 | 5.29E-10 | 16 |
| BORG5 | 52.45 | 5.71 | 1.12E-51 | 89 | 50.70 | 5.66 | 9.19E-51 | 85 |
| PIK3R1 | 1.75 | 0.81 | | 5 | 0.99 | -0.01 | | 5 |
| PAR-6G | 8.42 | 3.07 | 2.16E-05 | 16 | 9.75 | 3.28 | 2.82E-05 | 15 |
| DAAM1 | 4.59 | 2.20 | 3.60E-07 | 38 | 4.72 | 2.24 | 2.21E-07 | 37 |
| SPEC1 | 17.01 | 4.09 | 7.61E-04 | 15 | 8.68 | 3.12 | 1.93E-02 | 14 |
| ACK1 | 6.53 | 2.71 | 3.42E-07 | 25 | 6.20 | 2.63 | 7.87E-06 | 20 |
| BORG2 | 0.57 | -0.80 | 6.15E-01 | 7 | 0.78 | -0.35 | 8.14E-01 | 7 |
| PAK1 | 2.73 | 1.45 | 4.99E-03 | 15 | 2.41 | 1.27 | 3.00E-02 | 14 |
| PAK2 | 11.53 | 3.53 | 4.93E-13 | 49 | 10.22 | 3.35 | 9.03E-13 | 48 |
| FMNL2 | 4.46 | 2.16 | 2.37E-04 | 15 | 7.68 | 2.94 | 9.27E-06 | 16 |
| MRCKβ | 15.90 | 3.99 | 1.75E-19 | 101 | 27.12 | 4.76 | 7.28E-25 | 99 |
| CIP4 | 4.37 | 2.13 | 4.16E-01 | 8 | 5.49 | 2.46 | 2.49E-01 | 10 |
| IRSp53 | 12.60 | 3.65 | 3.55E-09 | 33 | 5.99 | 2.58 | 9.63E-07 | 31 |
| MRCKα | 13.11 | 3.71 | 7.51E-60 | 212 | 13.50 | 3.75 | 1.01E-52 | 208 |
| N-WASP | 25.36 | 4.66 | 1.19E-21 | 58 | 44.89 | 5.49 | 1.49E-25 | 54 |
| PAR-6B | 13.30 | 3.73 | 1.93E-17 | 56 | 16.47 | 4.04 | 5.87E-19 | 53 |
| BORG1 | 64.10 | 6.00 | 3.06E-20 | 41 | 70.40 | 6.14 | 5.99E-16 | 35 |
| PAK4 | 36.67 | 5.20 | 9.07E-38 | 64 | 52.29 | 5.71 | 1.38E-42 | 59 |
| BORG4 | 188.52 | 7.56 | 1.04E-11 | 22 | 242.06 | 7.92 | 1.85E-12 | 22 |

B

| GEFs | | | | | | | | |
|---|---|---|---|---|---|---|---|---|
| Protein | Cdc42b-CA/GFP | | | | Cdc42u-CA/GFP | | | |
| | Ratio | Log2 | p-value | Peptides used | Ratio | Log2 | p-value | Peptides used |
| α-PIX | 1.23 | 0.29 | 8.59E-01 | 8 | 2.26 | 1.18 | 2.66E-02 | 9 |
| RhoGEF | 0.54 | -0.88 | 1.21E-01 | 8 | 0.41 | -1.27 | 7.52E-03 | 12 |
| DOCK7 | 0.41 | -1.29 | 2.27E-32 | 184 | 0.50 | -1.01 | 2.78E-25 | 179 |
| DOCK6 | 0.96 | -0.06 | 9.11E-01 | 14 | 1.31 | 0.39 | 2.37E-01 | 13 |
| DOCK9 | 1.02 | 0.02 | 9.59E-01 | 13 | 1.07 | 0.09 | 8.84E-01 | 11 |
| β-PIX | 2.03 | 1.02 | 2.46E-02 | 17 | 1.96 | 0.97 | 4.14E-01 | 11 |
| Tuba | 7.89 | 2.98 | 8.00E-22 | 119 | 9.44 | 3.24 | 2.74E-22 | 117 |

C

| GAPs | | | | | | | | |
|---|---|---|---|---|---|---|---|---|
| Proteins | Cdc42b-CA/GFP | | | | Cdc42u-CA/GFP | | | |
| | Ratio | Log2 | p-value | Peptides used | Ratio | Log2 | p-value | Peptides used |
| RICH1 | 2.07 | 1.05 | 5.84E-04 | 29 | 1.00 | 0.01 | 9.90E-01 | 27 |
| DYNLL1 | 0.22 | -2.22 | 1.28E-05 | 16 | 0.22 | -2.19 | 3.39E-06 | 16 |
| RACGAP1 | 0.70 | -0.52 | 7.51E-03 | 28 | 0.64 | -0.64 | 3.12E-03 | 30 |
| STARD13 | 1.04 | 0.06 | 9.50E-01 | 8 | 1.04 | 0.06 | 8.24E-01 | 9 |
| DLC1 | 8.86 | 3.15 | 3.07E-03 | 8 | 7.85 | 2.97 | 1.02E-03 | 8 |
| RhoGAP31 | 1000.00 | 9.97 | | 3 | 1000.00 | 9.97 | | 4 |
